## Supplementary material for "Role of multiple pericentromeric repeats on heterochromatin assembly": Revised Supplemental Information

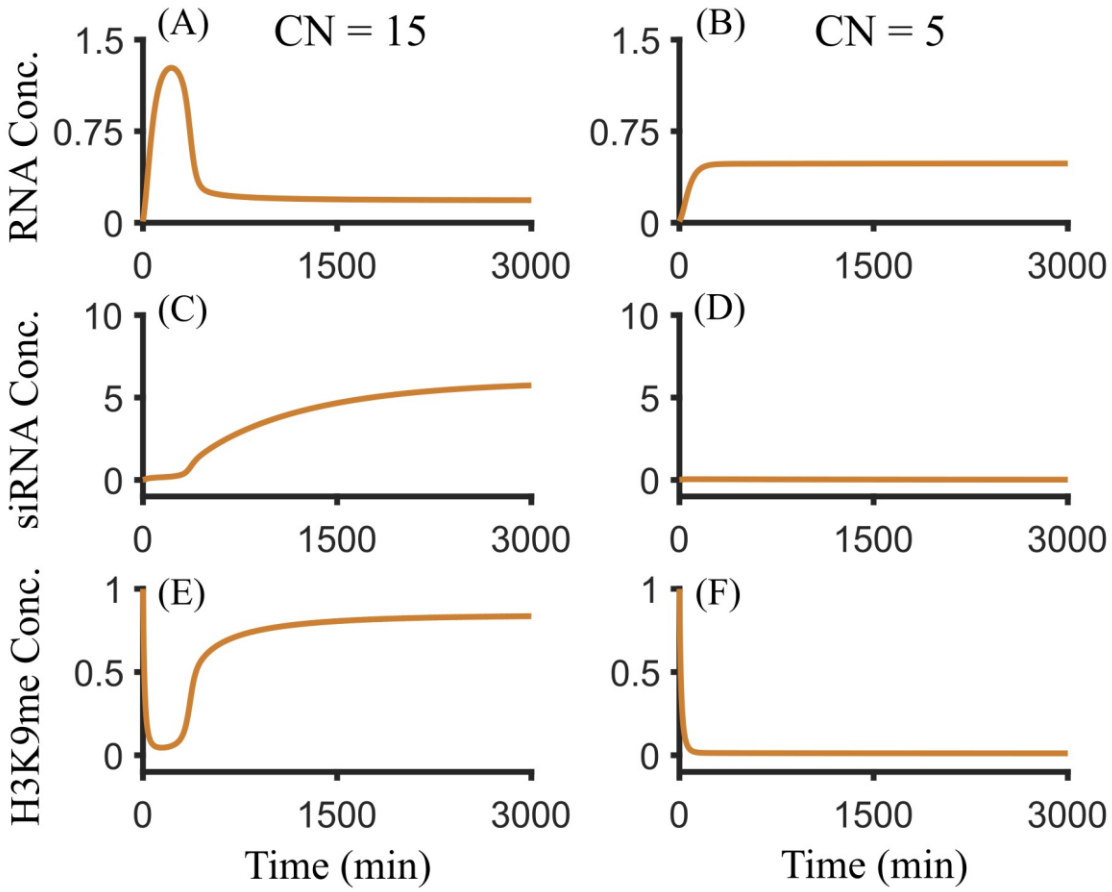


**Fig. S1.** ODE solution of eqs.1-3 for (A, C, E) CN = 15 and (B, D, F) CN = 5 for the same initial condition. (A, C, E) When CN = 15, the system is silenced, which is represented by high steady-state H3K9me concentration. (B, D, F) when CN = 5, the system is desilenced, which is represented by low steady-state H3K9me concentration.

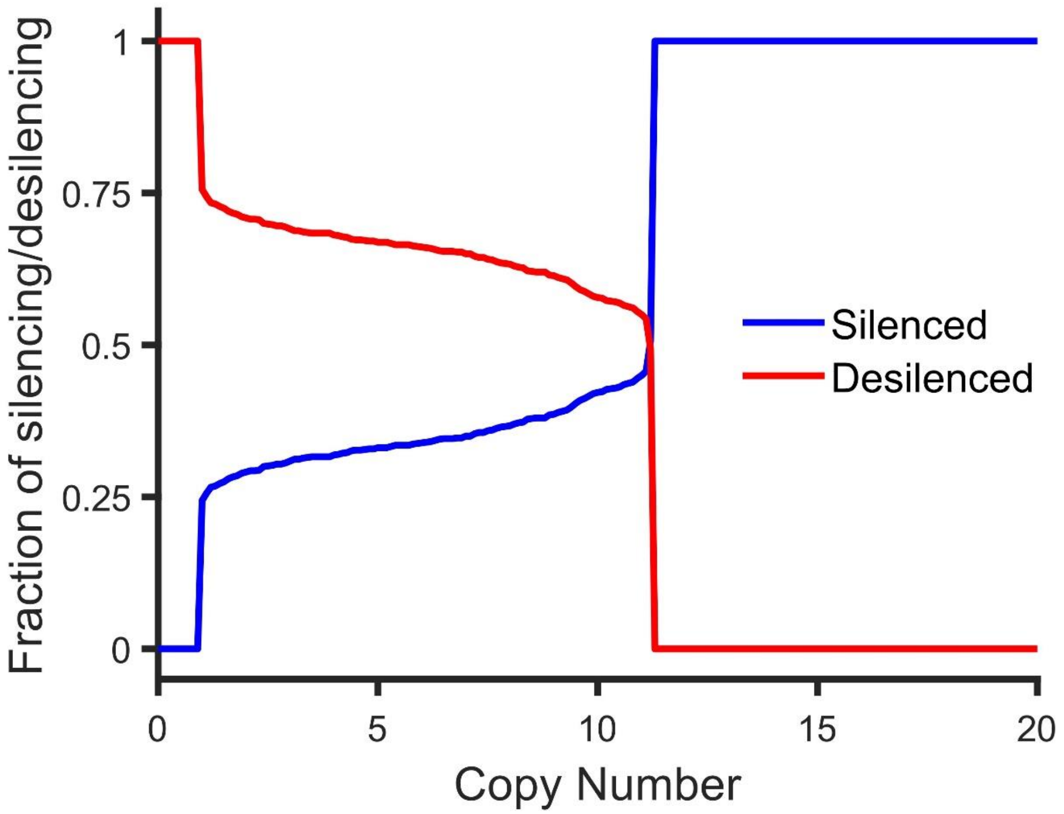


**Fig. S2.** Fraction of silenced and desilenced states obtained for 1000 different initial conditions. Bistable region is from CN=1 to CN=11.

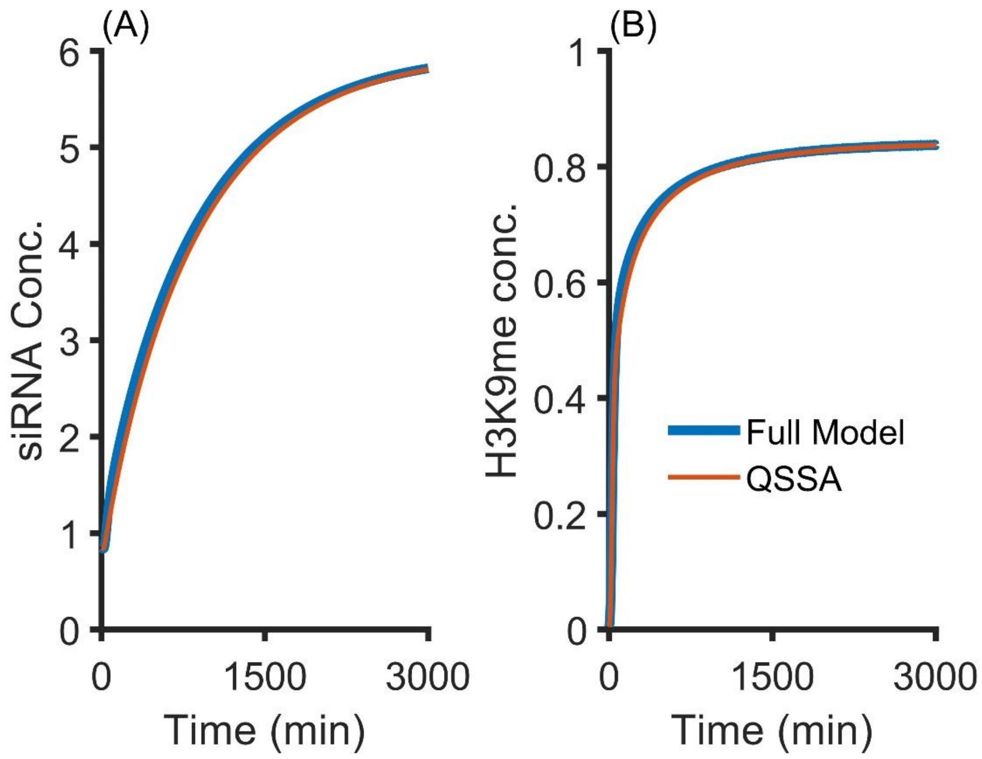


**Fig. S3.** Comparison of time evolutions of (A) siRNA, and (B) H3K9me between full model and QSSA at CN=15. Both siRNA (A) and (B) H3K9me concentrations evolve at similar rates between the full model and QSSA.

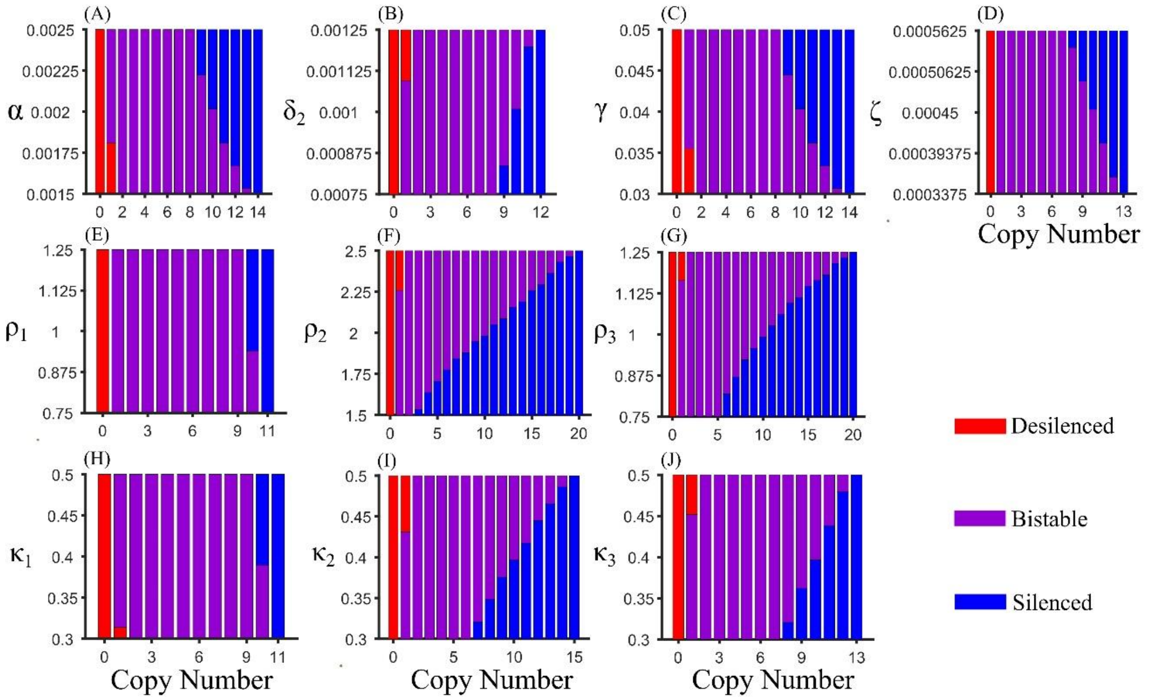


**Fig. S4:** Range of parameters (within 25% of their default values) facilitating desilenced, silenced, or bistable states at different copy numbers. Other than one parameter, all other parameters are held at the default values. High values of (A) $\alpha$, (C) $\gamma$, (D) $\zeta$, (E) $\rho_1$, and (H) $\kappa_1$ favor silencing, whereas high values of (B) $\delta_2$, (F) $\rho_2$, (G) $\rho_3$, (I) $\kappa_2$, and (J) $\kappa_3$ favor bistability or desilencing. $\alpha$ = Transcription Rate, $\delta_2$ = siRNA degradation rate, $\gamma$ = siRNA biogenesis rate, $\zeta$ = Basal methylation rate, $\rho_1$, $\rho_2$, and $\rho_3$ = Hill coefficient for transcription, siRNA biogenesis, methylation by siRNA respectively. $\kappa_1$,$\kappa_2$, and $\kappa_3$ = Half maximum methylation for transcription, siRNA biogenesis, and methylation by siRNA respectively.

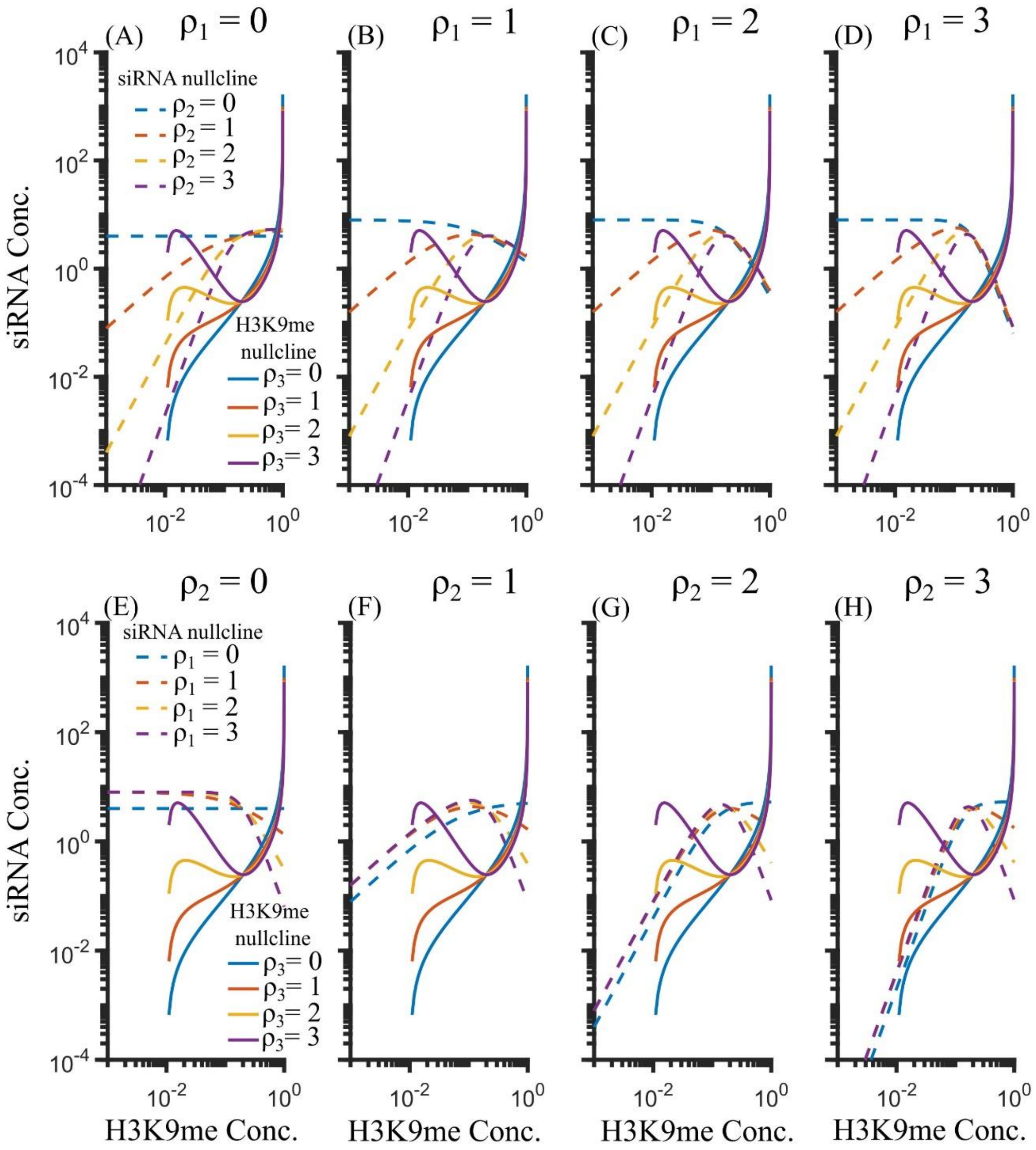


**Fig. S5.** Hill coefficient dependence of nullclines. H3K9me nullcline and siRNA nullcline can intersect at three distinct points if $\rho_1 \geq 1$, $\rho_2 \geq 2$, and $\rho_3 \geq 1$ in the reference parameters.

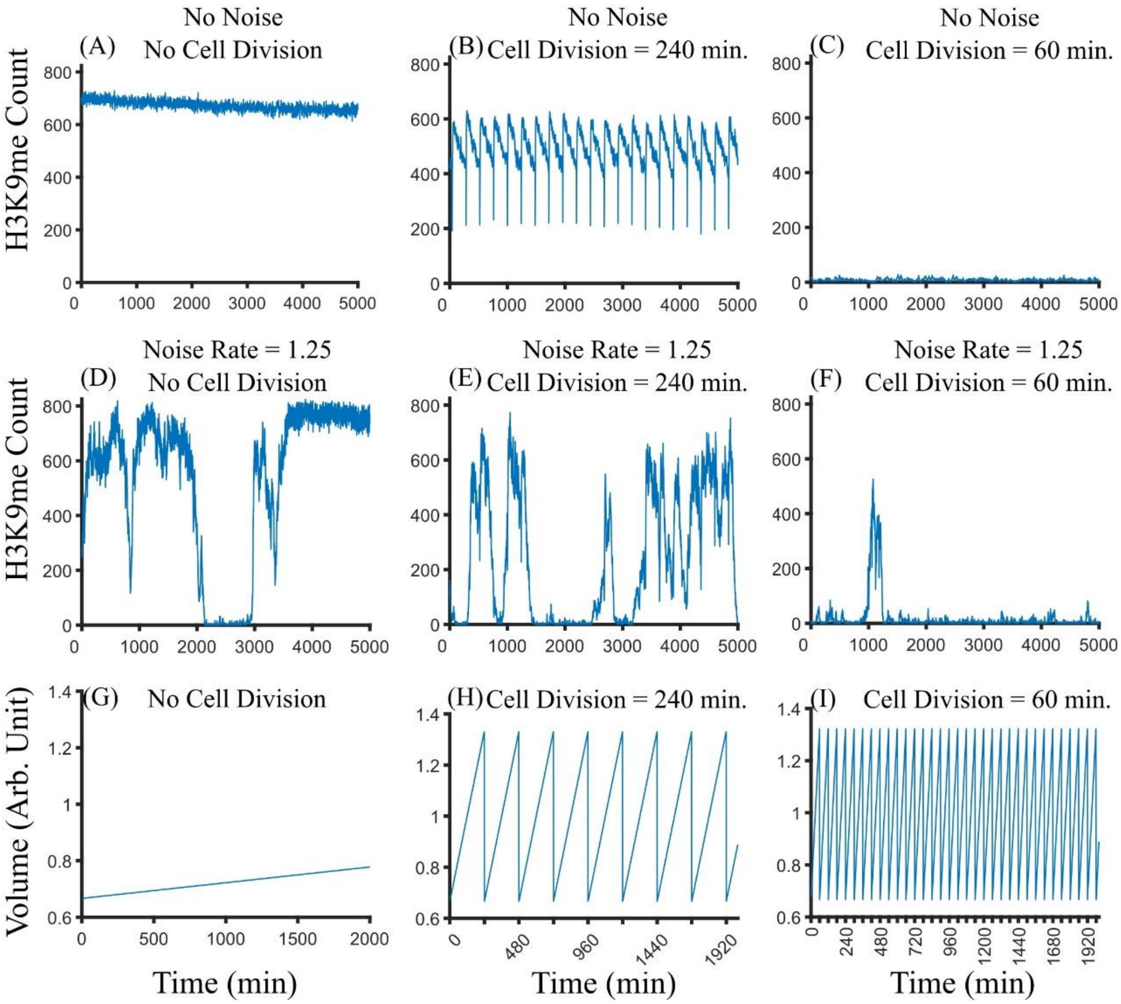


**Fig. S6.** Stochastic trajectories for copy number 15 for indicated noise level and cell division. (A) Silenced state is favored for the system with no cell division and noise. (B-C) Cell division can lead to desilencing. (D-F) The H3K9me profile with noise. (G-I), the volume growth with respect to time, for different cell division of the system.

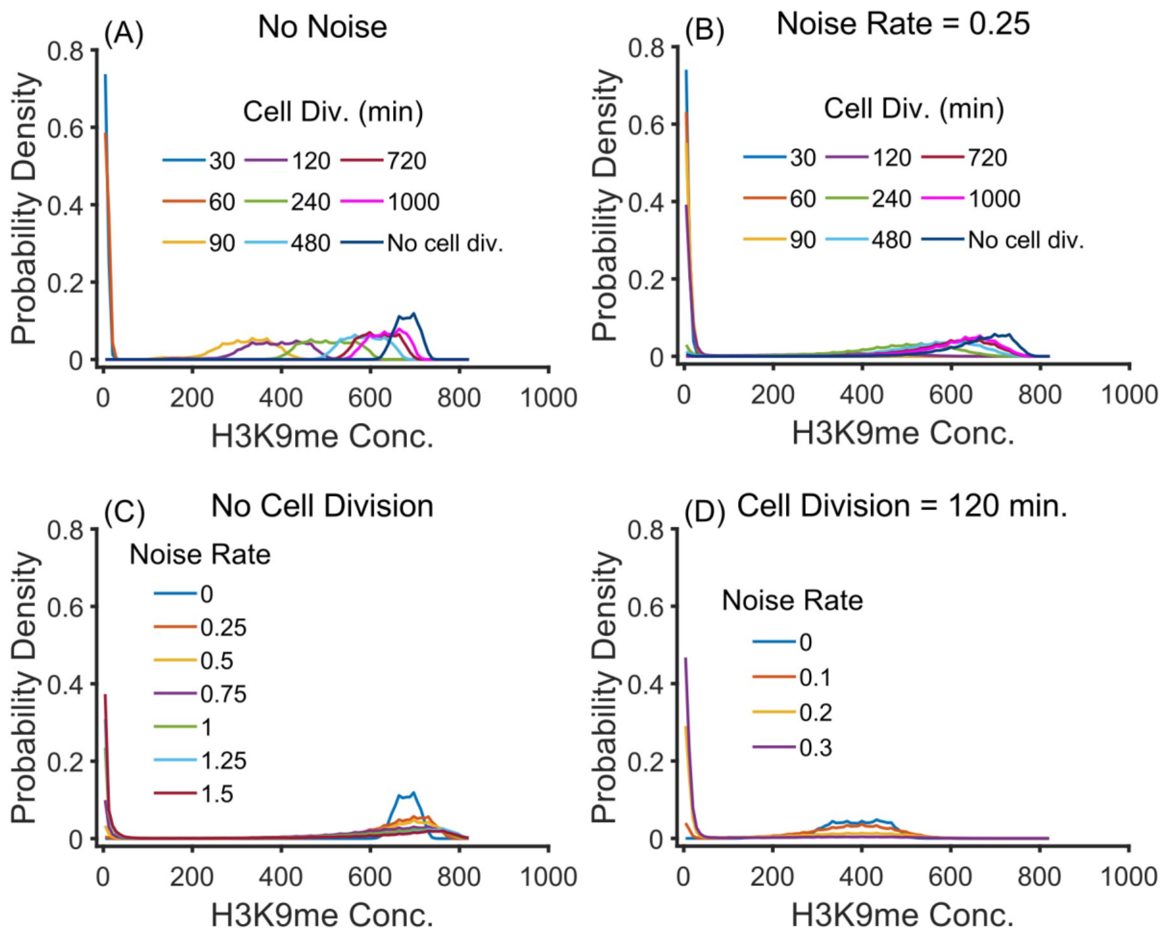


**Fig. S7.** The probability density of H3K9me concentration over time at indicated noise rate and cell division for copy number 15. Faster cell division favors the desilenced state for a system (A) without noise and (B) with noise. Increase in noise brings the system down to desilenced state (C) without cell division and (D) with cell division.

**Table S1.** Parameters identified by our analysis that facilitate silencing or desilencing.

| **Silencing** | **Desilencing** |
|---|---|
| Transcription rate ($\alpha$) | Half maximum methyl concentration for siRNA biogenesis ($\kappa_2$) |
| SiRNA biogenesis rate ($\gamma$) | Hill coefficient for siRNA biogenesis ($\rho_2$) |
| Methylation rate ($\epsilon$) | SiRNA degradation rate ($\delta_2$) |
| Methylation spreading rate ($\phi$) | Demethylation rate ($\delta_3$) |
| Basal methylation rate ($\zeta$) | RNA degradation rate ($\delta_1$) |
| Half maximum methylation conc. for transcription ($\kappa_1$) | Half maximum methylation conc. for methylation ($\kappa_3$) |
| Hill coefficient for transcription ($\rho_1$) | Hill coefficient for methylation ($\rho_3$) |

**Table S2.** Parameters used for solving differential eqs. (1-3) for WT cell

| Parameters | Values for WT cells |
|---|---|
| Transcription rate constant ($\alpha$) | 0.002 $mol. s^{-1}$ |
| RNA degradation rate constant ($\delta_1$) | 0.02 $s^{-1}$ |
| siRNA biogenesis rate ($\gamma$) | 0.04 $s^{-1}$ |
| Half maximum methylation conc. for transcription ($\kappa_1$) | 0.4 $mol$ |
| Hill coefficient for transcription ($\rho_1$) | 1 |
| Half maximum methylation conc. for siRNA biogenesis ($\kappa_2$) | 0.4 mol |
| Hill coefficient for siRNA biogenesis ($\rho_2$) | 2 |
| Half maximum methylation conc. for methylation ($\kappa_3$) | 0.4 mol |
| Hill coefficient for methylation ($\rho_3$) | 1 |
| siRNA degradation rate constant ($\delta_2$) | 0.001 $s^{-1}$ |
| Methylation rate constant ($\epsilon$) | 0.1 $mol^{-1}s^{-1}$ |
| Demethylation rate constant ($\delta_3$) | 0.083 $s^{-1}$ |
| Methylation spreading rate constant ($\phi$) | 0.04 $mol^{-1}s^{-1}$ |
| Basal methylation rate constant ($\zeta$) | 0.00045 $s^{-1}$ |

**Table S3.** Initial conditions applied in the simulation. Methylation concentration is normalized to 1.

| S.N | Initial condition (RNA, siRNA, H3K9me). Unit (mol) |
|---|---|
| 1 | (1, 1, 0) |
| 2 | (0.1, 0.1, 0.9) |
| 3 | (0, 1, 1) |
| 4 | (0.01, 0.01, 0.01) |
| 5 | (0, 0.5, 0) |
| 6 | (0.5, 0.5, 0.5) |
| 7 | (0.85, 1.5, 1) |
| 8 | (0.75, 0.35, 1) |
| 9 | (1.9, 2, 1) |
| 10 | (0, 3, 1) |
| 11 | (0.01, 0.01, 0.02) |
| 12 | (0.01, 0.01, 0.05) |
| 13 | (0.01, 0.01, 0.5) |
| 14 | (0.01, 0.01, 1) |
| 15 | (0.5, 0.01, 1) |
| 16 | (0.5, 0.1, 1) |
| 17 | (1,2,1) |
| 18 | (0.4, 0.6, 0) |
| 19 | (0, 0, 0.1) |
| 20 | (0.1, 0, 0) |
| 21 | (0.75, 0, 0.75) |
| 22 | (0, 0, 1) |
| 23 | (0, 4, 0) |
| 24 | (1,4,1) |
